## Supplementary Figures for "NiCo Identifies Extrinsic Drivers of Cell State Modulation by Niche Covariation Analysis"

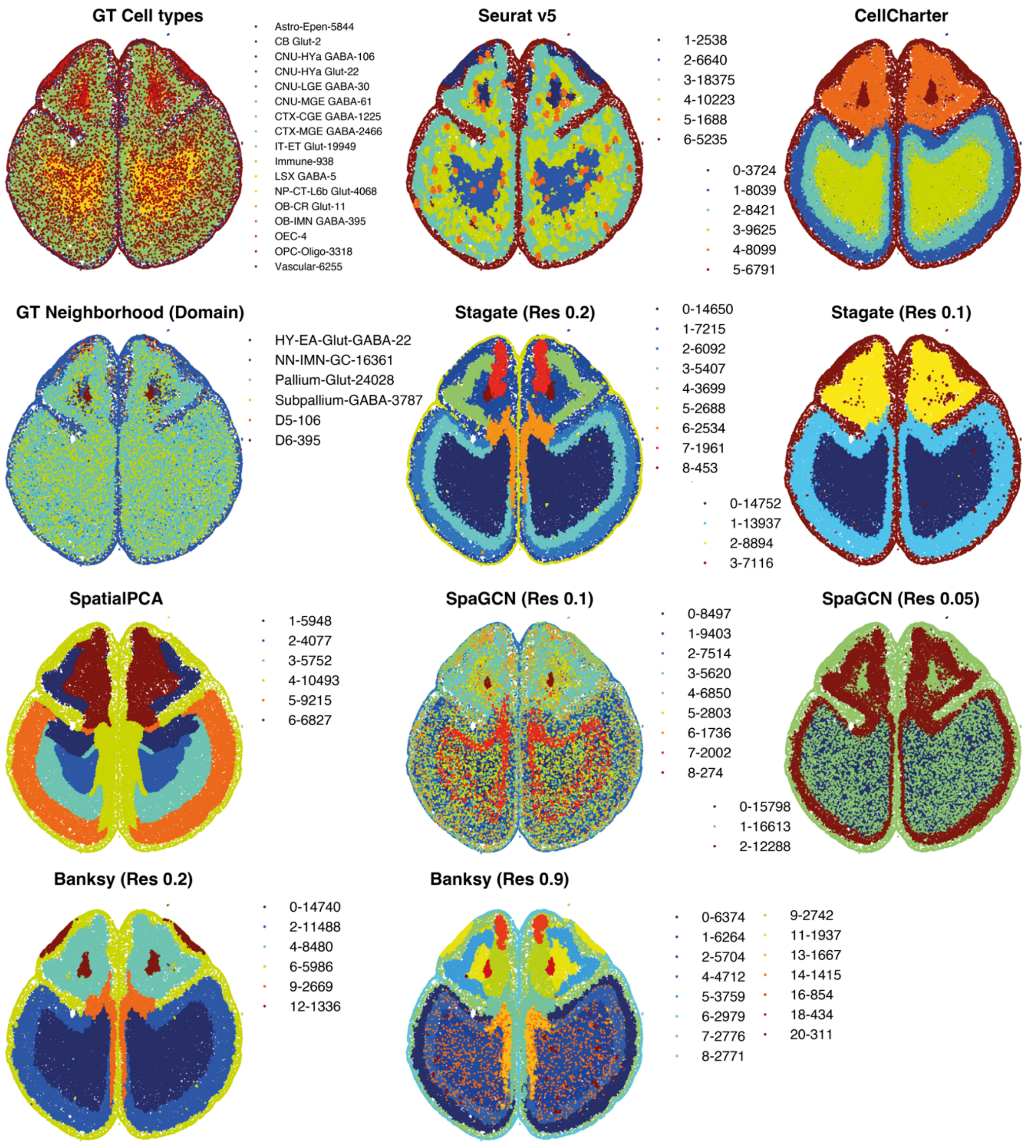


**Extended Data Fig. 1| Tissue domain detection in Allen Brain MERFISH Atlas data by different methods**. The spatial maps highlight topological domains on a brain section according to the authors' ground truth (GT) annotation, and predictions obtained by tailored niche detection methods as indicated. Some methods were run with different resolution parameters as indicated to match the number of GT domains (Methods).

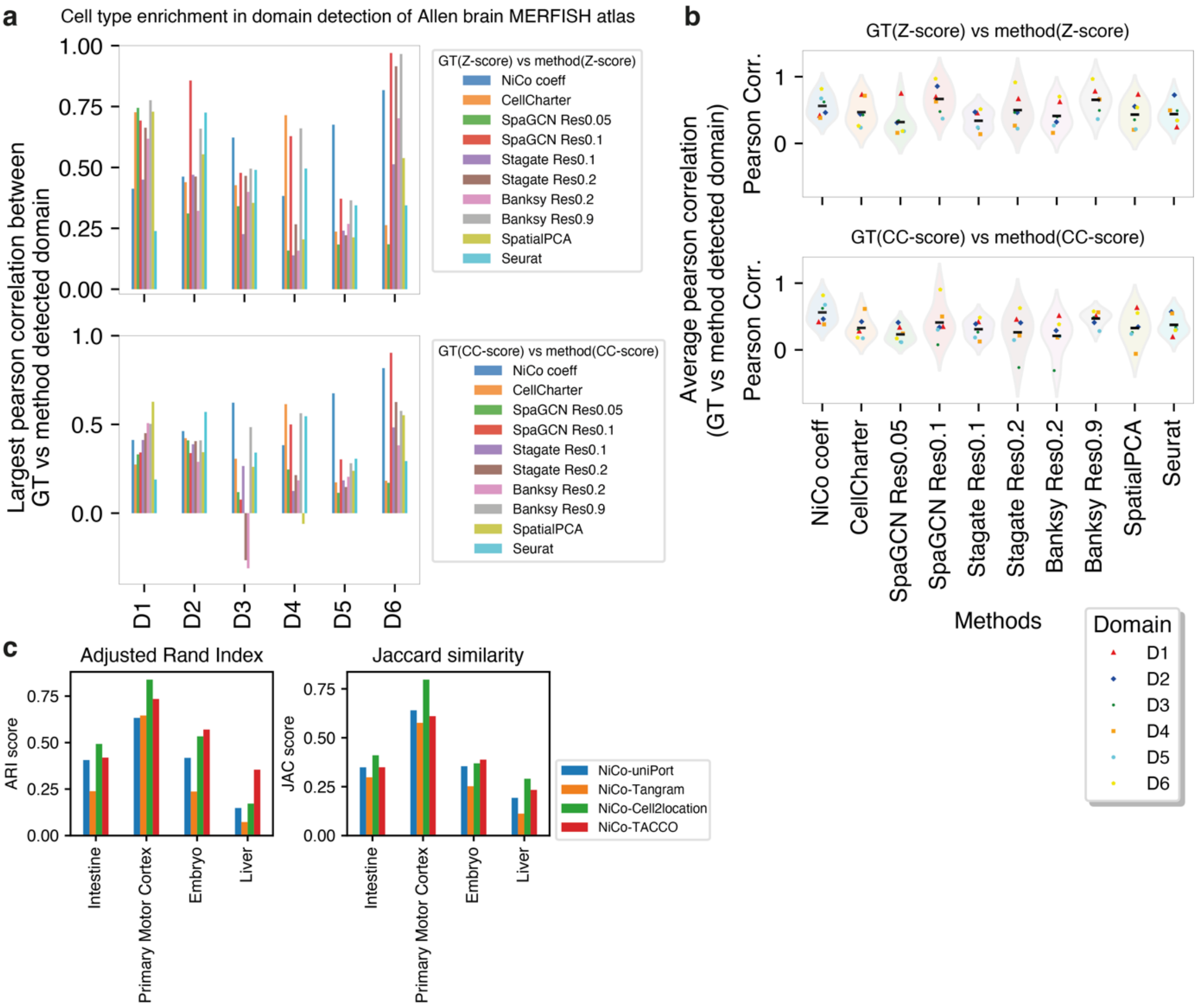


**Extended Data Fig. 2| Benchmarking of tissue domain detection on Allen brain MERFISH atlas data and cell type annotation**. (**a**, **b**), Cell type enrichment in GT domains was computed as enrichment Z-score (top) or CellCharter (CC) enrichment score (bottom). The Pearson correlation of this score is shown for the predicted domain with the highest correlation to the ground truth domain. For NiCo, the central cell type with the highest correlation was selected and the interaction coefficients were correlated to the Z-score/CC-score. Data are represented as barplots across domains (a) or violin plots across methods (b). The domain names are D1 (HY-EA-Glut-GABA), D2 (NN-IMN-GC), D3 (Pallium-Glut), D4 (Subpallium-GABA), D5 (D4; D1), D6 (D4; D2). Black line in (b), average correlation. **c**, comparison of NiCo annotations with uniPort, Tangram, TACCO, and cell2location for published datasets (mouse intestine and primary motor cortex MERFISH data, mouse embryo seqFISH data, and liver MERSCOPE data). Consistency of annotations between two methods was evaluated by Jaccard similarity (JAC) and adjusted rand index (ARI).

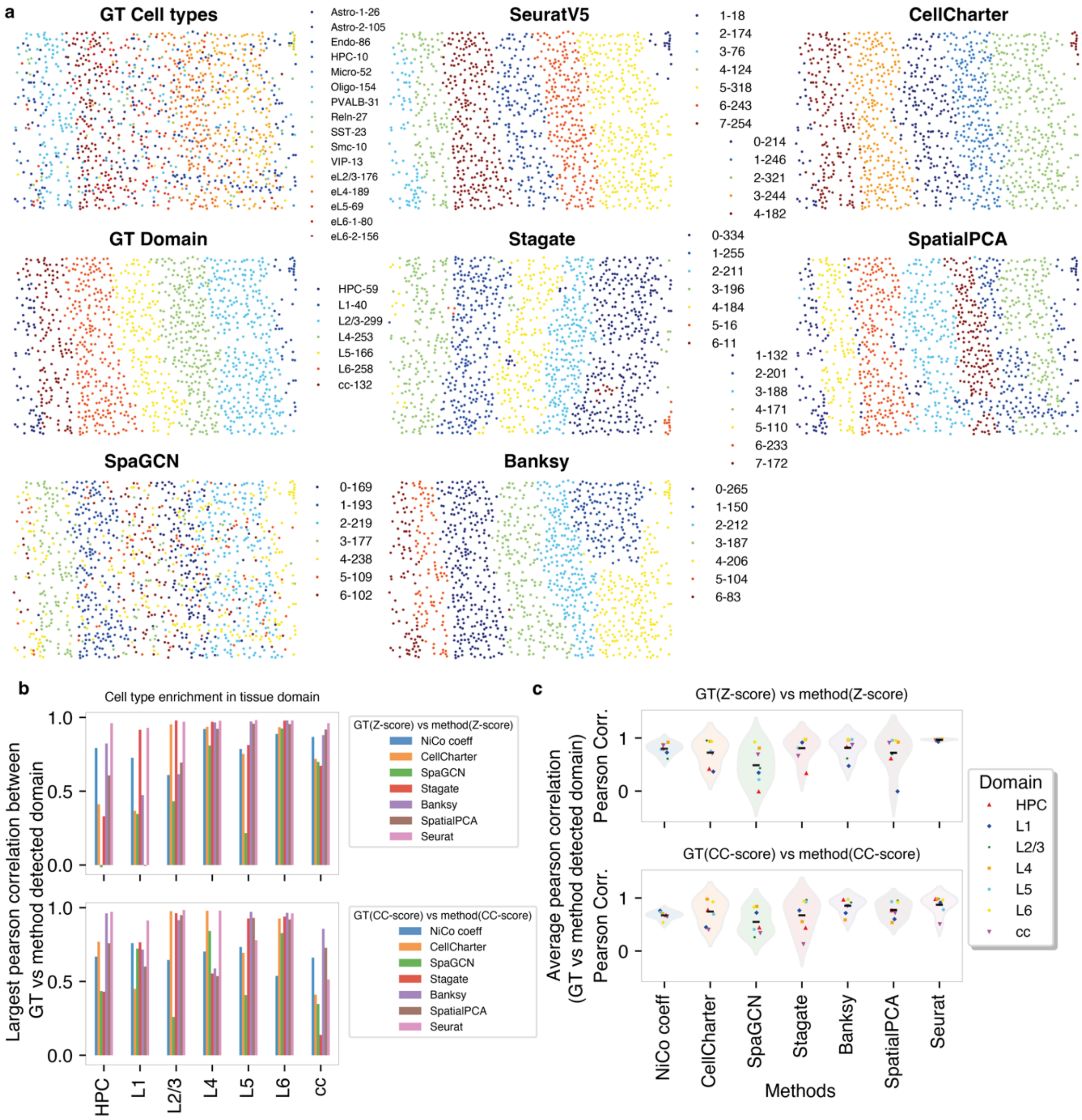


**Extended Data Fig. 3| Tissue domain detection on STARmap visual cortex data. a**, Predicted niche domains are highlighted in spatial maps of the STARmap visual cortex data for GT annotation, Seurat V5, CellCharter, Stagate, SpatialPCA, SpaGCN, and Banksy. Domain names are defined as L1, L2/3, L4, L5, L6, HPC (hippocampus) and cc (corpus callosum). (**b**, **c**), Cell type enrichment in GT domains was computed as enrichment Z-score (top) or CellCharter (CC) enrichment score (bottom). The Pearson correlation of this score is shown for the predicted domain with the highest correlation to the ground truth domain. For NiCo, the central cell type with the highest correlation was selected and the interaction coefficients were correlated to the Z-score/CC-score. Data are represented as barplots across domains (b) or violin plots across methods (c). Black line in (c), average correlation.

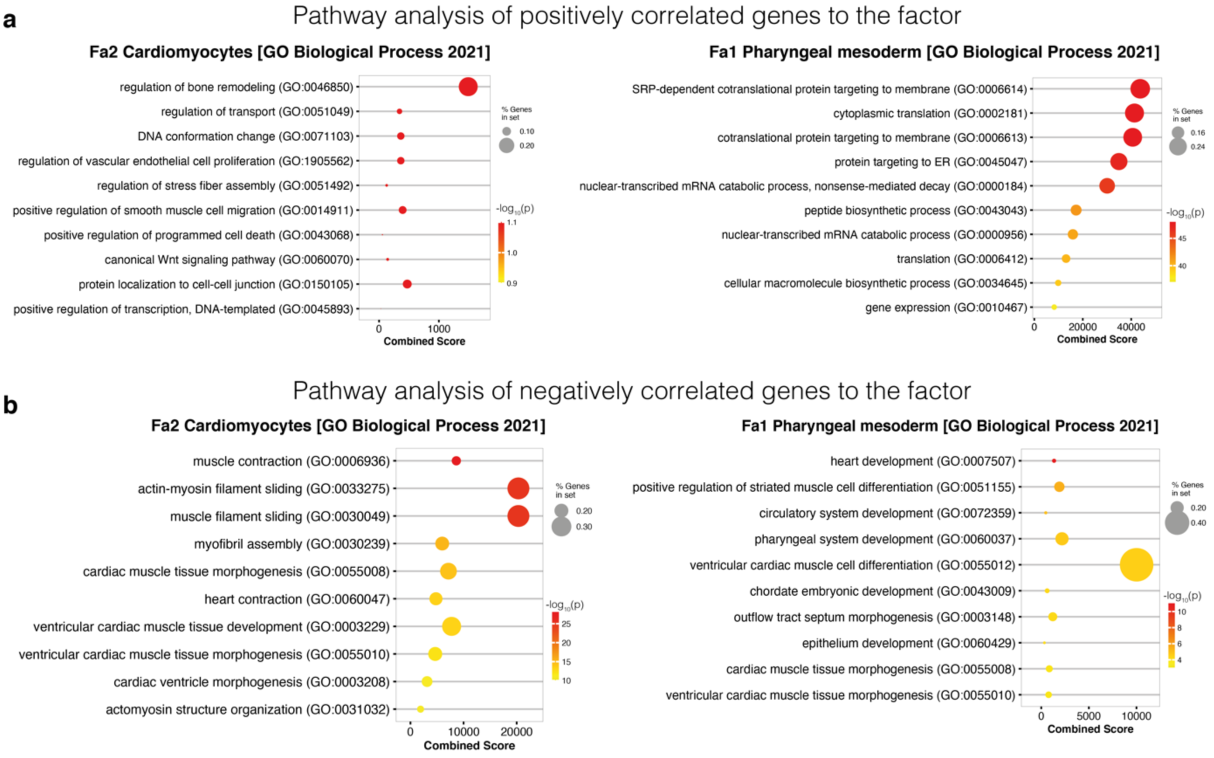


**Extended Data Fig. 4| Pathway analysis of factors associated with cardiomyocytes and pharyngeal mesoderm in mouse embryonic data. a,** The top 50 positively correlated genes, identified by Spearman correlation with cardiomyocytes Fa2 (left) and pharyngeal mesoderm Fa1 (right) were analyzed for Gene Ontology (GO) Biological Process enrichment. **b,** The top 50 negatively correlated genes, identified by Spearman correlation with cardiomyocytes Fa2 (left) and pharyngeal mesoderm Fa1 (right) were analyzed for Gene Ontology (GO) Biological Process enrichment. See Methods for details.

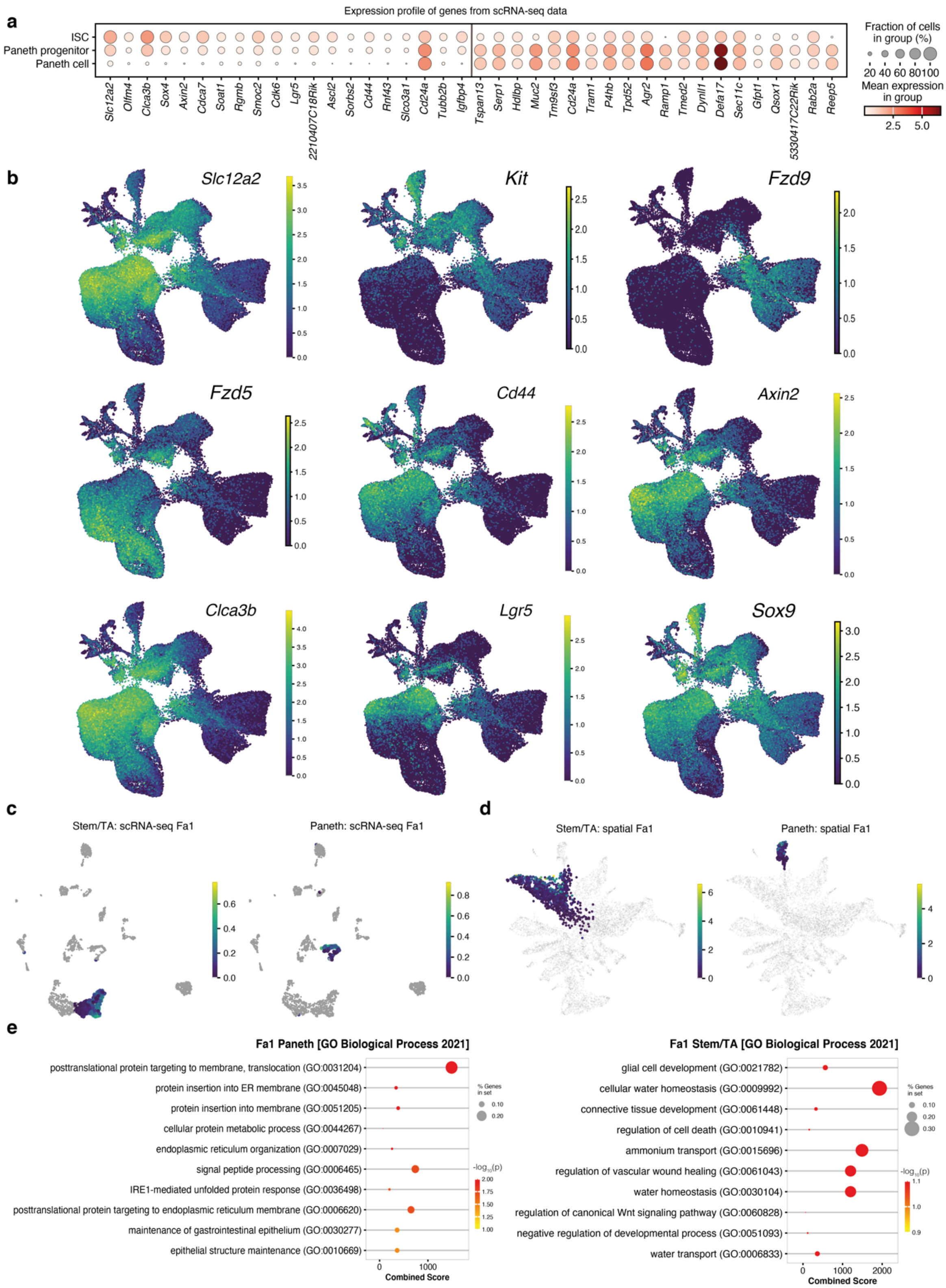


**Extended Data Fig. 5| Pathway analysis and expression of genes associated with Stem/TA and Paneth cell covariation in mouse intestine. a,** The top 20 positively correlated genes to Paneth Fa1 (left) and stem/TA Fa1 (right) are displayed as dot plot highlighting the fraction of cells in the respective population expressing a gene (dot size) and the mean expression level (dot color). **b,** Selected genes upregulated in intestinal stem cells and Paneth cell progenitors are highlighted in a UMAP representation of mouse intestinal scRNA-seq data from Böttcher et al.[53]. **(c-d),** Stem/TA Fa1 and Paneth Fa1 are highlighted in the a UMAP representation of the scRNA-seq reference data [54] (c) and the spatial data [31] (d). **e,** The top 50 positively correlated genes, identified by Spearman correlation with Paneth Fa2 (left) and Stem/TA Fa1 (right) were analyzed for Gene Ontology (GO) Biological Process enrichment. See Methods for details.

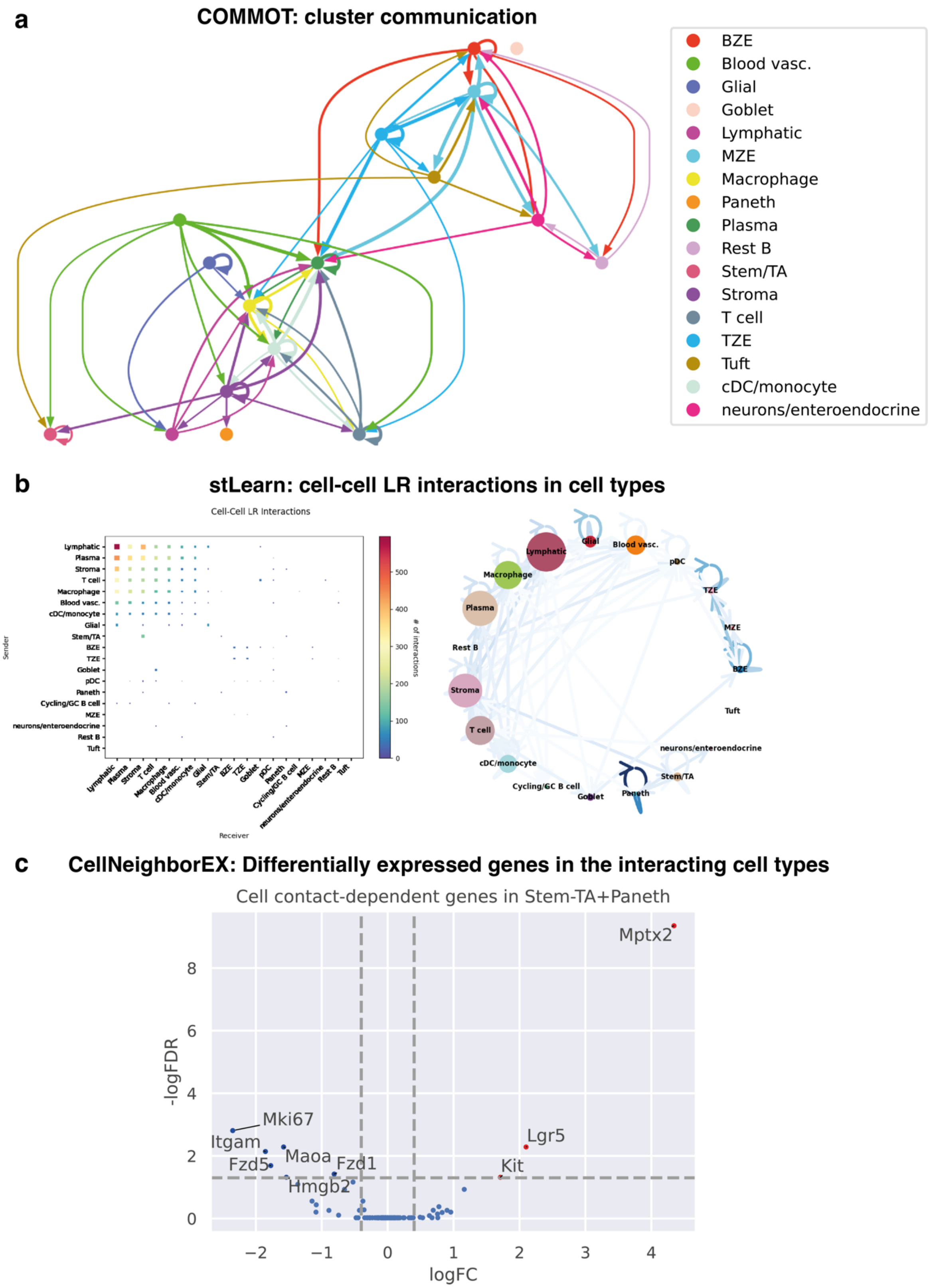


**Extended Data Fig. 6| Benchmarking on cell-cell interaction analysis in the intestinal dataset. a**, The cluster communication graph for pathways (total-total) from COMMOT[21]. **b**, Cell type-cell type LR interactions from stLearn[57]. **c**, Contact-dependent genes between stem/TA and Paneth cells from CellNeighborEx[58].

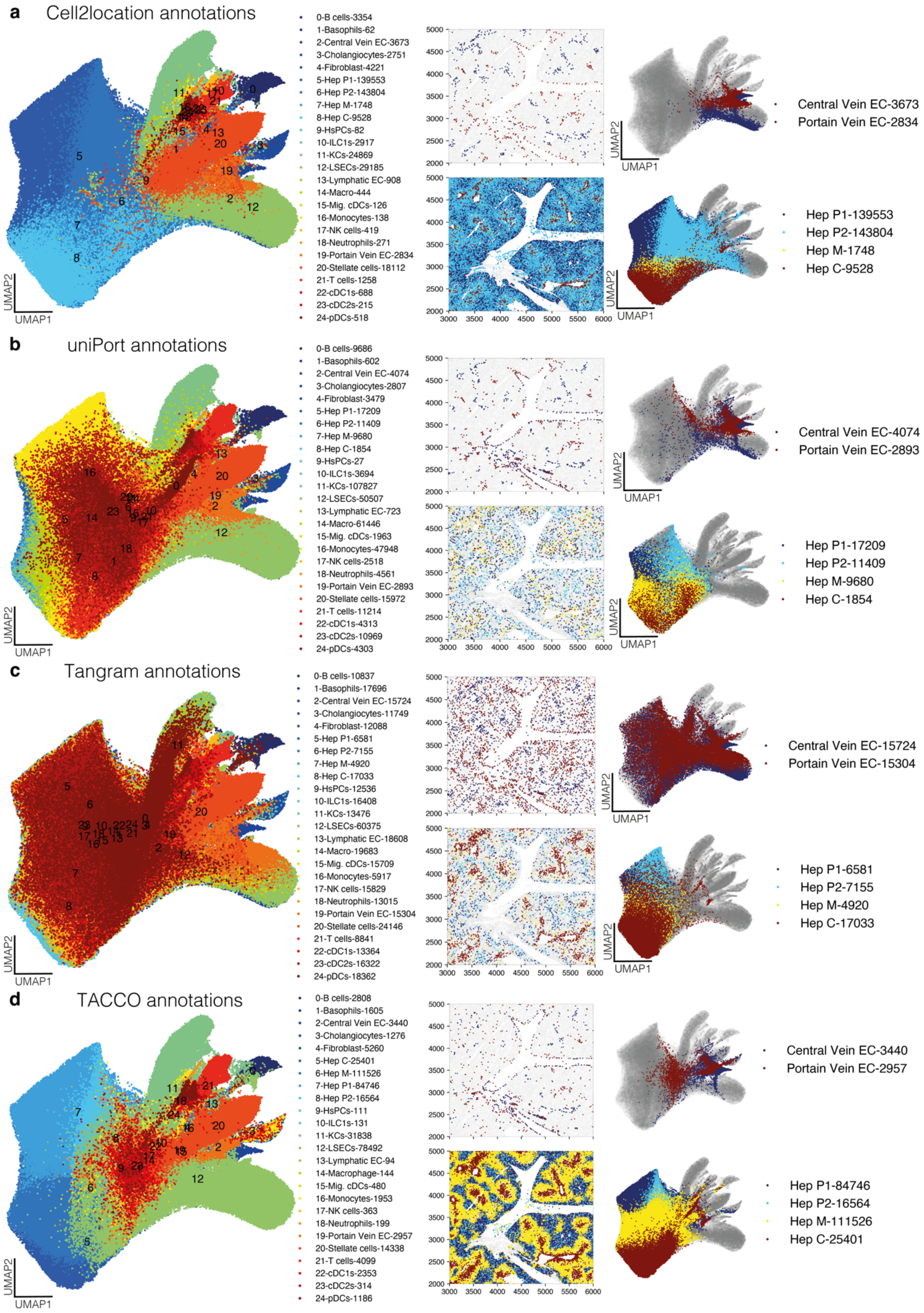

a
Cell2location annotations
0-B cells-3354
1-Basophils-62
2-Central Vein EC-3673
3-Cholangiocytes-2751
4-Fibroblast-4221
5-Hep P1-139553
6-Hep P2-143804
7-Hep M-1748
8-Hep C-9528
9-HsPCs-82
10-ILC1s-2917
11-KCs-24869
12-LSECs-29185
13-Lymphatic EC-908
14-Macro-444
15-Mig. cDCs-126
16-Monocytes-138
17-NK cells-419
18-Neutrophils-271
19-Portain Vein EC-2834
20-Stellate cells-18112
21-T cells-1258
22-cDC1s-688
23-cDC2s-215
24-pDCs-518
Central Vein EC-3673
Portain Vein EC-2834
Hep P1-139553
Hep P2-143804
Hep M-1748
Hep C-9528
UMAP1
UMAP2
b
uniPort annotations
0-B cells-9686
1-Basophils-602
2-Central Vein EC-4074
3-Cholangiocytes-2807
4-Fibroblast-3479
5-Hep P1-17209
6-Hep P2-11409
7-Hep M-9680
8-Hep C-1854
9-HsPCs-27
10-ILC1s-3694
11-KCs-107827
12-LSECs-50507
13-Lymphatic EC-723
14-Macro-61446
15-Mig. cDCs-1963
16-Monocytes-47948
17-NK cells-2518
18-Neutrophils-4561
19-Portain Vein EC-2893
20-Stellate cells-15972
21-T cells-11214
22-cDC1s-4313
23-cDC2s-10969
24-pDCs-4303
Central Vein EC-4074
Portain Vein EC-2893
Hep P1-17209
Hep P2-11409
Hep M-9680
Hep C-1854
c
Tangram annotations
0-B cells-10837
1-Basophils-17696
2-Central Vein EC-15724
3-Cholangiocytes-11749
4-Fibroblast-12088
5-Hep P1-6581
6-Hep P2-7155
7-Hep M-4920
8-Hep C-17033
9-HsPCs-12536
10-ILC1s-16408
11-KCs-13476
12-LSECs-60375
13-Lymphatic EC-18608
14-Macro-19683
15-Mig. cDCs-15709
16-Monocytes-5917
17-NK cells-15829
18-Neutrophils-13015
19-Portain Vein EC-15304
20-Stellate cells-24146
21-T cells-8841
22-cDC1s-13364
23-cDC2s-16322
24-pDCs-18362
Central Vein EC-15724
Portain Vein EC-15304
Hep P1-6581
Hep P2-7155
Hep M-4920
Hep C-17033
d
TACCO annotations
0-B cells-2808
1-Basophils-1605
2-Central Vein EC-3440
3-Cholangiocytes-1276
4-Fibroblast-5260
5-Hep C-25401
6-Hep M-111526
7-Hep P1-84746
8-Hep P2-16564
9-HsPCs-111
10-ILC1s-131
11-KCs-31838
12-LSECs-78492
13-Lymphatic EC-94
14-Macrophage-144
15-Mig. cDCs-480
16-Monocytes-1953
17-NK cells-363
18-Neutrophils-199
19-Portain Vein EC-2957
20-Stellate cells-14338
21-T cells-4099
22-cDC1s-2353
23-cDC2s-314
24-pDCs-1186
Central Vein EC-3440
Portain Vein EC-2957
Hep P1-84746
Hep P2-16564
Hep M-111526
Hep C-25401

**Extended Data Fig. 7| Cell type annotation of spatial mouse liver data by alternative methods. (a-d),** Left: UMAP representation based on gene expression of the spatial data. Cell type annotations are highlighted. The legend indicates the number of cells for each cell type. Middle: Select cell types are highlighted for a representative tissue area. Top, central vein and portal vein endothelial cells (EC). Bottom, hepatocytes of different zones, i.e., central (Hep C), mid-zonal (Hep M) and portal (Hep P1/P2). Right: Cell types from (Middle) are highlighted in the expression UMAP representation. Results are shown for cell2location (a), uniPort (b), Tangram (c) and TACCO (d).

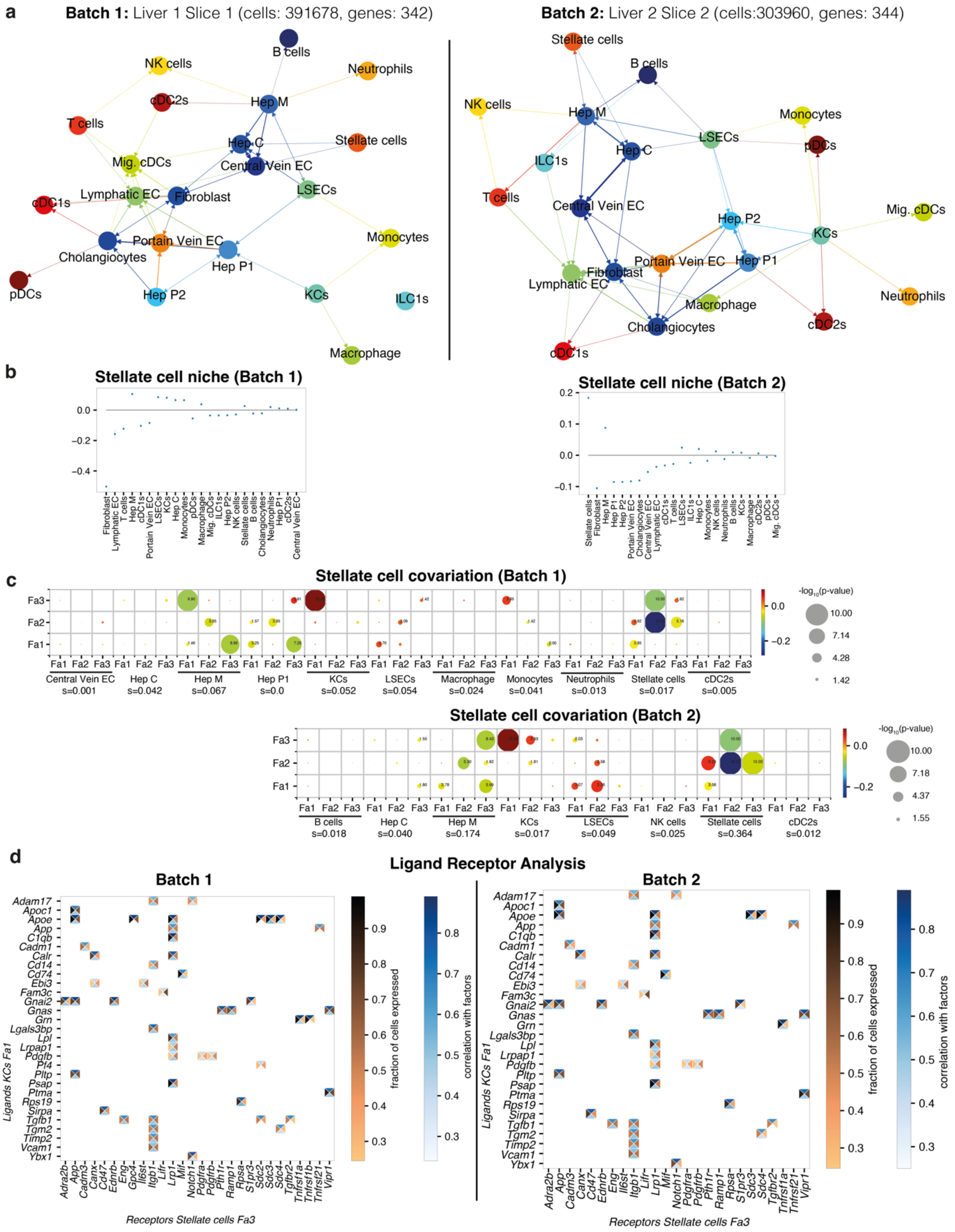


**Extended Data Fig. 8| NiCo analysis on two batches of the liver dataset.** NiCo was run on experimental batches from the Vizgen liver dataset, i.e., slice 1 of liver 1 (batch 1) and slice 2 of liver 2 (batch 2). **a**, Cell type interaction map (interaction threshold c= 0.1) for batch 1 (left) and batch 2 (right). **b**, The logistic regression coefficients of the stellate cell niche for batch 1 (left) and batch 2 (right). **c**, Covariation between stellate factors (y-axis) and co-localized neighborhood cell type factors (x-axis) for batch 1 (top) and batch 2 (bottom). Circle size scales linearly with $-\log_{10}$(p-value), and circle color indicates ridge regression coefficients. S denotes the normalized niche coefficient score. **d**, Ligand-receptor pairs correlated with stellate cell Fa3 and KC Fa1 for batch 1 (left) and batch 2 (right). See Methods. The rectangle's north and south faces represent ligand and receptor correlation to the factors, while west and east faces represent the proportion of ligand and receptor expressing cells.

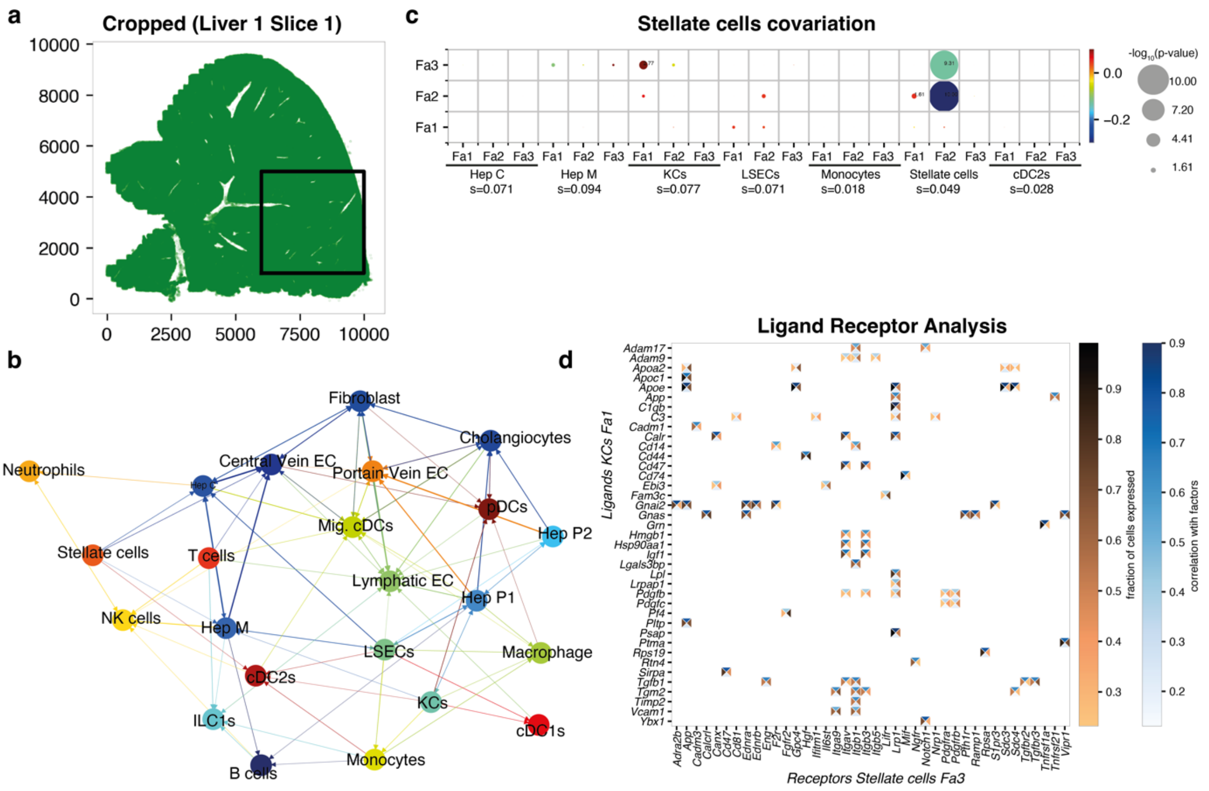


**Extended Data Fig. 9| NiCo analysis on cropped region of liver data. a,** Spatial map of slice 1 from the Vizgen liver dataset analyzed in Fig. 5. The cropped region analyzed with NiCo is indicated by a black box and contains a subset of 88,772 cells (22.7%). **b**, Cell-type niche interaction map (interaction threshold c=0.1). **c**, Covariation between stellate factors (y-axis) and co-localized neighborhood cell type factors (x-axis). Circle size scales linearly with -log$_{10}$(p-value), and circle color indicates ridge regression coefficients. S denotes the normalized niche coefficient score. **d**, Ligand-receptor pairs correlated with stellate cell Fa3 and KC Fa1. See Methods. The rectangle's north and south faces represent ligand and receptor correlation to the factors, while west and east faces represent the proportion of ligand and receptor expressing cells.

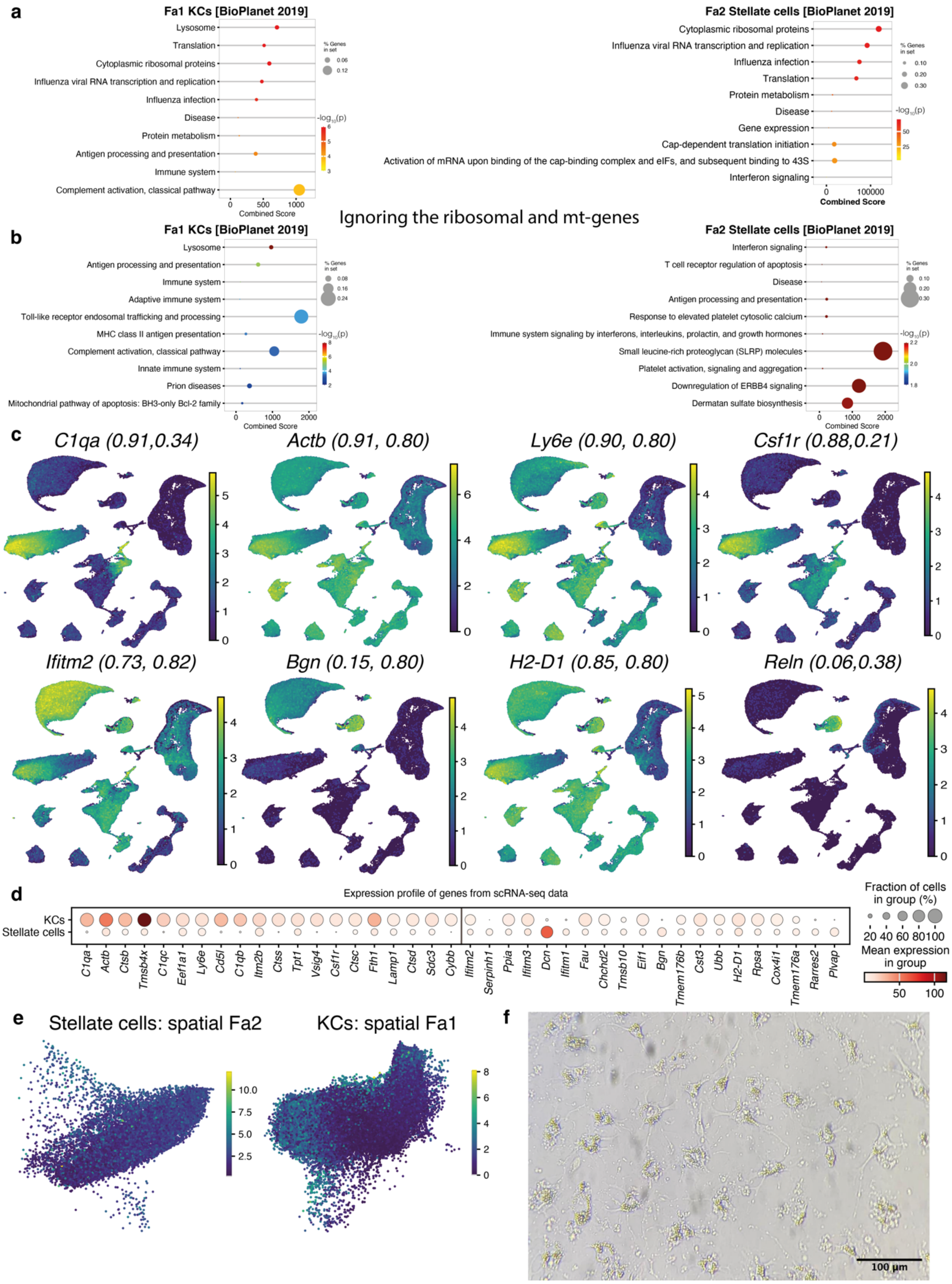


**Extended Data Fig. 10| Pathway analysis and expression of genes associated with Stellate cells and Kupffer cells covariation in mouse liver. a,** The top 50 positively correlated genes, identified by Spearman correlation with KCs Fa1 (left) and stellate cells Fa2 (right) were analyzed for the enrichment of gene sets from the "Bio Planet 2019" database. See Methods for details. **b,** Same as (a) but after removing genes encoding mitochondrial genes, and genes encoding ribosomal subunits. **c,** UMAP representation of scRNA-seq/snRNA-seq/CITE-seq reference data [63] highlighting normalized expression of select genes correlating to stellate cells Fa2 (top) or KCs Fa1. The Pearson correlation coefficients between the gene expression and the respective factor values (KC Fa1, stellate cells Fa2) are indicated in the parentheses. **d,** The top 20 positively correlated genes to KCs Fa1(left) and stellate cells Fa2 (right) are displayed as dot plot highlighting the fraction of cells in the respective population expressing a gene (dot size) and the mean expression level (dot color). **e,** UMAP representation of spatial data for stellate cells (left) and KCs (right) highlighting stellate cell Fa2 and KCs Fa1, respectively. Both factors exhibit expression gradients in the respective population. **f,** Brightfield image of hepatic stellate cell *in vitro culture* 48 hours after isolation.

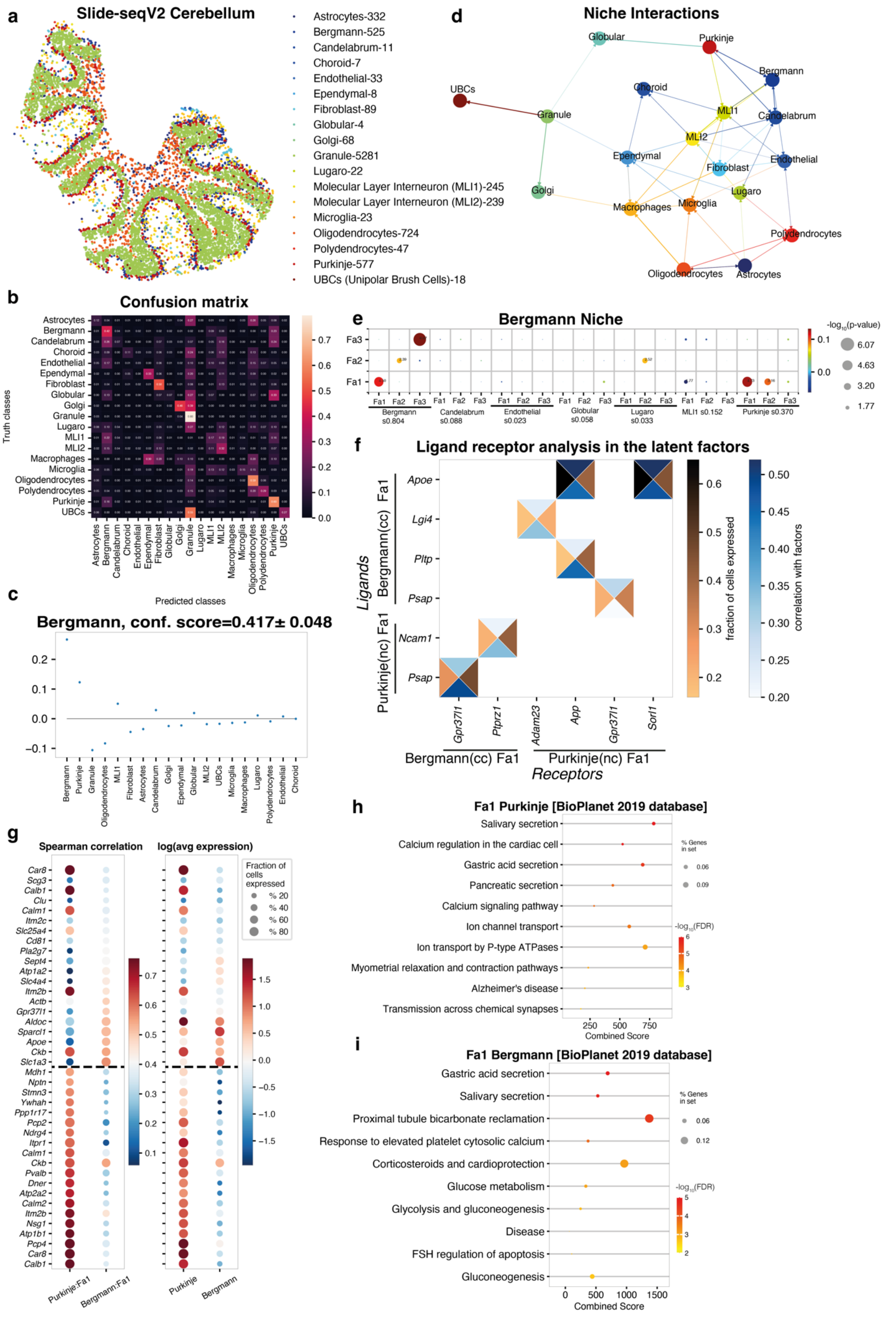
a
Slide-seqV2 Cerebellum
Astrocytes-332
Bergmann-525
Candelabrum-11
Choroid-7
Endothelial-33
Ependymal-8
Fibroblast-89
Globular-4
Golgi-68
Granule-5281
Lugaro-22
Molecular Layer Interneuron (MLI1)-245
Molecular Layer Interneuron (MLI2)-239
Microglia-23
Oligodendrocytes-724
Polydendrocytes-47
Purkinje-577
UBCs (Unipolar Brush Cells)-18
d
Niche Interactions
Globular
Purkinje
Bergmann
UBCs
Choroid
Granule
ML1
Candelabrum
ML2
Ependymal
Endothelial
Fibroblast
Golgi
Lugaro
Macrophages
Microglia
Polydendrocytes
Oligodendrocytes
Astrocytes
b
Confusion matrix
Truth classes
Predicted classes
Astrocytes
Bergmann
Candelabrum
Choroid
Endothelial
Ependymal
Fibroblast
Globular
Golgi
Granule
Lugaro
MLI1
MLI2
Macrophages
Microglia
Oligodendrocytes
Polydendrocytes
Purkinje
UBCs
e
Bergmann Niche
Fa1
Fa2
Fa3
Bergmann s0.804
Candelabrum s0.088
Endothelial s0.023
Globular s0.058
Lugaro s0.033
MLI1 s0.152
Purkinje s0.370
-log10(p-value)
6.07
4.63
3.20
1.77
c
Bergmann, conf. score=0.417± 0.048
Bergmann
Purkinje
Granule
Oligodendrocytes
MLI1
Fibroblast
Astrocytes
Candelabrum
Golgi
Ependymal
Globular
MLI2
UBCs
Microglia
Macrophages
Lugaro
Polydendrocytes
Endothelial
Choroid
f
Ligand receptor analysis in the latent factors
Ligands
Bergmann(cc) Fa1
Purkinje(nc) Fa1
Apoe
Lgi4
Pltp
Psap
Ncam1
Psap
Gpr37l1
Ptprz1
Adam23
App
Gpr37l1
Sorl1
Bergmann(cc) Fa1
Purkinje(nc) Fa1
Receptors
fraction of cells expressed
correlation with factors
g
Spearman correlation
log(avg expression)
Fraction of cells expressed
% 20
% 40
% 60
% 80
Car8
Scg3
Calb1
Clu
Calm1
Itm2c
Slc25a4
Cd81
Pla2g7
Sept4
Atp1a2
Slc4a4
Itm2b
Actb
Gpr37l1
Aldoc
Sparcl1
Apoe
Ckb
Slc1a3
Mdh1
Nptn
Stmn3
Ywhah
Ppp1r17
Pcp2
Ndrg4
Itpr1
Calm1
Ckb
Pvalb
Dner
Atp2a2
Calm2
Itm2b
Nsg1
Atp1b1
Pcp4
Car8
Calb1
Purkinje:Fa1
Bergmann:Fa1
Purkinje
Bergmann
h
Fa1 Purkinje [BioPlanet 2019 database]
Salivary secretion
Calcium regulation in the cardiac cell
Gastric acid secretion
Pancreatic secretion
Calcium signaling pathway
Ion channel transport
Ion transport by P-type ATPases
Myometrial relaxation and contraction pathways
Alzheimer's disease
Transmission across chemical synapses
% Genes in set
-log10(FDR)
Combined Score
i
Fa1 Bergmann [BioPlanet 2019 database]
Gastric acid secretion
Salivary secretion
Proximal tubule bicarbonate reclamation
Response to elevated platelet cytosolic calcium
Corticosteroids and cardioprotection
Glucose metabolism
Glycolysis and gluconeogenesis
Disease
FSH regulation of apoptosis
Gluconeogenesis
% Genes in set
-log10(FDR)
Combined Score

**Extended Data Fig. 11| NiCo application on Slide-seqV2 cerebellum data**. **a**, The spatial map of cell types in the cerebellum as annotated in ref. [74]. **b**, NiCo confusion matrix. **c**, The logistic regression coefficients of the Bergmann glia niche. **d**, The cell type niche interaction map (interaction threshold c=0.08). **e**, Covariation between Bergmann glia factors (y-axis) and co-localized neighborhood cell type factors (x-axis). Circle size scales linearly with $-\log_{10}$(p-value), and circle color indicates ridge regression coefficients. S denotes the normalized niche coefficient score. **f**, Ligand-receptor pairs correlated with Bergmann Fa1 and Purkinje Fa1. See Methods. The rectangle's north and south faces represent ligand and receptor correlation to the factors, while west and east faces represent the proportion of ligand and receptor expressing cells. cc, central cell; nc, niche cell. **g**, Spearman correlations (left) and average expression (right) for the top 20 positively correlated genes for Purkinje Fa1 and Bergmann Fa1. Dashed line demarcates the genes associated with each cell types. **h**, The top 50 positively correlated genes, identified by Spearman correlation with Purkinje Fa1 were analyzed for enriched pathways from the BioPlanet 2019 database. **i**, The top 50 positively correlated genes, identified by Spearman correlation with Bergmann Fa1 were analyzed for enriched pathways from the BioPlanet 2019 database.
